# Vicarious touch: overlapping neural patterns between seeing and feeling touch

**DOI:** 10.1101/2022.06.21.497107

**Authors:** S. Smit, D. Moerel, R. Zopf, A.N. Rich

## Abstract

Simulation theories propose that vicarious touch arises when seeing someone else being touched triggers corresponding representations of being touched. Prior electroencephalography (EEG) findings show that seeing touch modulates both early and late somatosensory responses (measured with or without direct tactile stimulation). Functional Magnetic Resonance Imaging (fMRI) studies have shown that seeing touch increases somatosensory cortical activation. These findings have been taken to suggest that when we see someone being touched, we simulate that touch in our sensory systems. The somatosensory overlap when seeing and feeling touch differs between individuals, potentially underpinning variation in vicarious touch experiences. Increases in amplitude (EEG) or cerebral blood flow response (fMRI), however, are limited in that they cannot test for the information contained in the neural signal: seeing touch may not activate the same *information* as feeling touch. Here, we use time-resolved multivariate pattern analysis on whole-brain EEG data from people with and without vicarious touch experiences to test whether seen touch evokes overlapping neural representations with the first-hand experience of touch. Participants felt touch to the fingers (*tactile* trials) or watched carefully matched videos of touch to another person’s fingers (*visual* trials). In both groups, EEG was sufficiently sensitive to allow decoding of touch location (little finger vs. thumb) on *tactile* trials. However, only in individuals who reported feeling touch when watching videos of touch could a classifier trained on *tactile* trials distinguish touch location on *visual* trials. This demonstrates that, for people who experience vicarious touch, there is overlap in the information about touch location held in the neural patterns when seeing and feeling touch. The timecourse of this overlap implies that seeing touch evokes similar representations to *later stages* of tactile processing. Therefore, while simulation may underlie vicarious tactile sensations, our findings suggest this involves an abstracted representation of directly felt touch.

## Introduction

Simulation theories propose that we empathise with others by mapping the observed state onto a similar representation within our own system (Bird and Viding, 2014; Gallese et al., 2004; Gazzola et al., 2006; Keysers and Gazzola, 2009). Such co-activation might apply not only to observed actions and emotions but also to sensations (Gallese, 2005; Keysers and Gazzola, 2009). For instance, sometimes people report that seeing others being touched results in similar tactile sensations on their own bodies. These vicarious touch (also referred to as ‘mirror-touch’) experiences could therefore rely on evoking similar neural representations when we see someone touched as when we feel touch directly (Bufalari and Ionta, 2013; Keysers and Gazzola, 2009).

Electroencephalography (EEG) and magnetoencephalography (MEG) studies show that seeing touch can modulate both early (45ms) and later (up to 700ms) somatosensory activity (Adler et al., 2016; Adler and Gillmeister, 2019; Bufalari et al., 2007; Galilee and McCleery, 2016; Martínez-Jauand et al., 2012; Pihko et al., 2010; Rigato et al., 2019b, 2019a; Streltsova and McCleery, 2014). For instance, several studies simultaneously presented visual stimuli showing touch to a body part (e.g., the hand) with tactile stimulation of the participant’s own corresponding body part. Seeing the touch, compared to seeing a no-touch stimulus (e.g., touch next to a hand), modulated very early (∼45ms; P45) somatosensory evoked potentials measured from electrodes centred on the somatosensory cortex (Adler et al., 2016; Adler and Gillmeister, 2019; Bufalari et al., 2007; Martínez-Jauand et al., 2012; Rigato et al., 2019b). As the P45 is likely to reflect activity in the primary somatosensory cortex (Allison et al., 1992; Schubert et al., 2008), it has been inferred that this modulation reflects a match between the representations of the seen and felt touch at *early sensory-perceptual* processing stages (Adler et al., 2016; Adler and Gillmeister, 2019; Bufalari et al., 2007; Martínez-Jauand et al., 2012; Rigato et al., 2019b), which is consistent with simulation theories (Gallese, 2005).

Studies using functional Magnetic Resonance Imaging (fMRI) report that seeing another person being touched evokes activity in brain regions typically responsive to the first-hand experience of touch (Blakemore et al., 2005; Ebisch et al., 2008; Holle et al., 2013; Keysers et al., 2004; Kuehn et al., 2014, 2013, 2018; Lee Masson et al., 2018; Schaefer et al., 2009, 2006). For example, Keysers et al., (2004) were the first to demonstrate that seeing brush strokes to another person’s legs caused similar activation in the secondary somatosensory cortex as having one’s own legs stroked. Subsequent studies have further shown that seeing touch can also activate the primary somatosensory cortex (Blakemore et al., 2005; Ebisch et al., 2008; Holle et al., 2013; Kuehn et al., 2014, 2013, 2018; Schaefer et al., 2009; but see Chan and Baker, 2015). Based on these findings, Keysers and colleagues (2009, 2010, 2022) propose that somatosensory activation in response to observing touch could underlie our ability to understand and empathise with others’ tactile experiences.

There is individual variability in the degree to which the somatosensory cortex is activated by seen touch. For instance, mirror-touch synaesthetes, who consistently feel touch on their own body when they see someone being touched, show stronger somatosensory activation when viewing touch compared to non-synaesthetic controls (Blakemore et al., 2005; Holle et al., 2013). A recent ultra-high field fMRI study in typical adults showed that simply viewing touch to another person’s fingers can activate fine-grained single finger maps in area 3b of the primary somatosensory cortex similar to directly feeling touch (Kuehn et al., 2018). However, not all participants showed this visually evoked somatosensory effect, and for those who did, some showed almost perfect overlap between tactile-and visually-driven activation whereas others had non-overlapping tactile/visual activations. Further, people vary substantially in their reports of how often and intensely they experience vicarious touch (for a review see Gillmeister et al., 2017). One possibility is that the degree to which visual and tactile neural representations of touch overlap could link to subjective differences in how we perceive touch to others.

The strong prediction of simulation theories is that the same *information* is activated when we observe others being touched as when we feel touch directly (Gallese, 2006, 2005). However, activation in the somatosensory cortex in response to seeing and feeling touch does not necessarily indicate that the same neural processes are driving this activation (Lamm and Majdandžić, 2015). Methods like fMRI and M/EEG provide a broad measure of neural activity, encompassing thousands of neurons. These neuron populations may exhibit diverse patterns under different conditions, but their overall metabolic activity may be similar, resulting in comparable hemodynamic responses or electrophysiological output. For example, seeing touch to a *little finger* versus a *thumb* is likely to result in a very similar event-related potential (ERP) measured over the somatosensory cortex. In addition, it is difficult to establish using ERPs whether brain activity evoked by *seeing* touch to a little finger is at all similar to that evoked by directly *feeling* touch to the little finger (as opposed to the thumb), as the input received via the tactile and visual modality is very different. Consequently, employing traditional univariate analyses that measure average *activation* are limited in evaluating the presence of shared representations and the information contained in the overlapping activation (Haynes and Rees, 2006; Hebart and Baker, 2018; Lamm and Majdandžić, 2015). Multivariate pattern analyses (MVPA or ‘decoding’ methods) take the *pattern of activation* across many sensors. According to simulation theories, seeing touch to different locations should elicit different activation patterns, analogous to those evoked by the first-hand experience of touch to those locations. This method allows us to directly compare the response evoked when one is touched versus observing touch to another, which gets much closer to testing the prediction from simulation theories (Lamm and Majdandžić, 2015).

In addition, MVPA uses directionless summary measures (e.g., classification accuracy) to quantify the dissimilarity between two conditions in each participant, regardless of the direction of the effect. In contrast, to be observed at the group level, traditional univariate analyses require consistency in the direction of neural effects across individuals. As previous fMRI findings show large inter-individual differences in somatosensory activation patterns when seeing and feeling touch to different fingers (Kuehn et al., 2018), MVPA seems a useful avenue for investigating shared representations between the modalities and the potential link to subjective experiences. Together, univariate and multivariate analyses complement each other in studying vicarious tactile processing.

The goal of the present study is to provide a more detailed understanding of the dynamic processing underlying vicarious touch experiences by testing if there is overlap in neural activation patterns when seeing and feeling touch (for the preregistration see: https://osf.io/9ed24/). To learn more about the presence of information^1^ over time, we apply multivariate analysis to EEG data, which takes the dynamic activation patterns across *many* EEG sensors covering the brain (Grootswagers et al., 2017). We specifically focus on decoding information about the location of touch on the body, as previous research suggests that observing touch elicits precise somatotopic representations in the tactile domain (Blakemore et al., 2005; Kammers et al., 2009; Riemer et al., 2014; Thomas et al., 2006). For our main analysis, we train a machine learning classifier at each timepoint to distinguish neural activation patterns when participants are touched on the little finger versus thumb (*tactile* trials) and test it on data when they *see* touch to the little finger versus thumb (*visual* trials; *between-modality* decoding). Our visual and tactile stimuli are carefully matched and controlled so that the classifier can only rely on information regarding the location of touch. If the classifier performs above chance, we can conclude that at those timepoints there is overlap in the neural activation patterns, suggesting the *shared information* in the neural signal between seeing and feeling touch includes the location of touch. We also test the tactile classifier on independent (left-out) *tactile* trials (*within-modality* decoding) to identify *when* the location of a felt touch is reflected in the neural pattern for comparison with the timecourse of any *between-modality* decoding.

We separate our sample into individuals with and without vicarious touch experiences to explore the degree to which any overlap in neural activation patterns (measured across the whole brain) between seeing and feeling touch is linked to the subjective experience of vicarious touch. We define these groups based on whether participants reported tactile experiences while watching videos of different types of touch (Smit et al., in prep). MVPA allows us to analyse each participant separately by training and testing the classifier on neural activation patterns specific to that individual. We then calculate average classification accuracies at each timepoint for each group (note that EEG signals are not averaged, just the classification accuracies).

The use of time-resolved EEG decoding allows us to test *when* any overlap between seeing and feeling touch occurs, which can tell us more about the underlying mechanisms. Simulation theories propose that seeing touch evokes a *sensory* representation of being touched (Gallese, 2005; Gallese and Sinigaglia, 2011). This could occur due to ‘direct matching’ of the seen touch with corresponding representations of the felt touch (e.g., Iacoboni et al., 1999; Rizzolatti et al., 2001; Rizzolatti and Sinigaglia, 2016). Alternatively, it could be due to predictive processes: seeing a touch results in the prediction that one will feel touch; the resulting prediction error (when no touch occurs) is ‘explained away’ by activating somatosensory cortex (Bach and Schenke, 2017; Ishida et al., 2015). The timecourse of overlap between the modalities will be informative about the level at which vicarious tactile processes occur, as overlap with an *early* tactile signal suggests the activation of a more *sensory* representation whereas overlap with a late tactile signal suggests a more *abstracted* representation.

## Materials and Methods

### Participants

We recruited right-handed naïve undergraduate students through Macquarie University participant pools. All participants reported normal or corrected to normal vision and no neurological disorders. As preregistered, we tested 20 participants before we performed statistical group-level Bayesian analyses. We then used our stopping rule and continued data collection until we had sufficient evidence for either the alternative-or null hypothesis for at least 50% of the timepoints (ranging from -1100ms to 1800ms with a resolution of ∼4ms) for the main analyses (within-and between-modality decoding of touched finger; see below). We interpreted a Bayes Factor (BF) of 6 (or 1/6 in case of evidence for the null) as sufficient evidence (Dienes, 2011; Wetzels et al., 2011). We tested 40 participants in total; six participants were excluded due to their performance on the experimental task being below the pre-specified threshold of 60% correct. Our final sample after exclusions consisted of 34 participants and of these, 16 were classified as having vicarious touch sensations and 18 as not having vicarious touch sensations based on their responses to the post-experiment vicarious touch video questionnaire (see Table 1). Participants provided written consent and either received $20 per hour or course credit in return for participation. The study was approved by the Macquarie University Ethics Committee.

**Table 1.** Sample characteristics and results of Bayesian Mann-Whitney Independent Samples t-tests.

|  | **Vicarious touch group**<br>(n = 16)<br>Mean (SD) | **No vicarious touch group**<br>(n = 18)<br>Mean (SD) | **Comparison**<br>**BF<sub>10</sub>** |
| --- | --- | --- | --- |
| Age in years | 20.06 (2.72)<br>Range 17-28 | 22.56 (5.93)<br>Range 18-40 | 0.565 |
| Sex | 4 males; 25% | 7 males; 39% | N/A |
| Vicarious touch reported out of 10 videos | 5 (2.80)<br>Range 1-10 | 0 (0.0) | N/A |
| Emotional reactivity score | 8.25 (2.67) | 9.44 (1.34) | 0.126 |

### Stimuli

We recorded EEG data while participants felt direct tactile stimulation to their own hand alternating with seeing videos depicting touch to another person’s hand. On *tactile* trials, the little finger or thumb of the participant’s right hand was stimulated, and on *visual* trials, participants viewed videos on the screen of a right hand being touched on the little finger or thumb (Fig. 1a, b). To control for external touch-side (left vs. right), both the participant’s own hand and the hand on the screen flipped between each run (palm up or palm down) so that touch to each individual finger occurred equally often on the left and the right of the screen.

**Figure 1.**
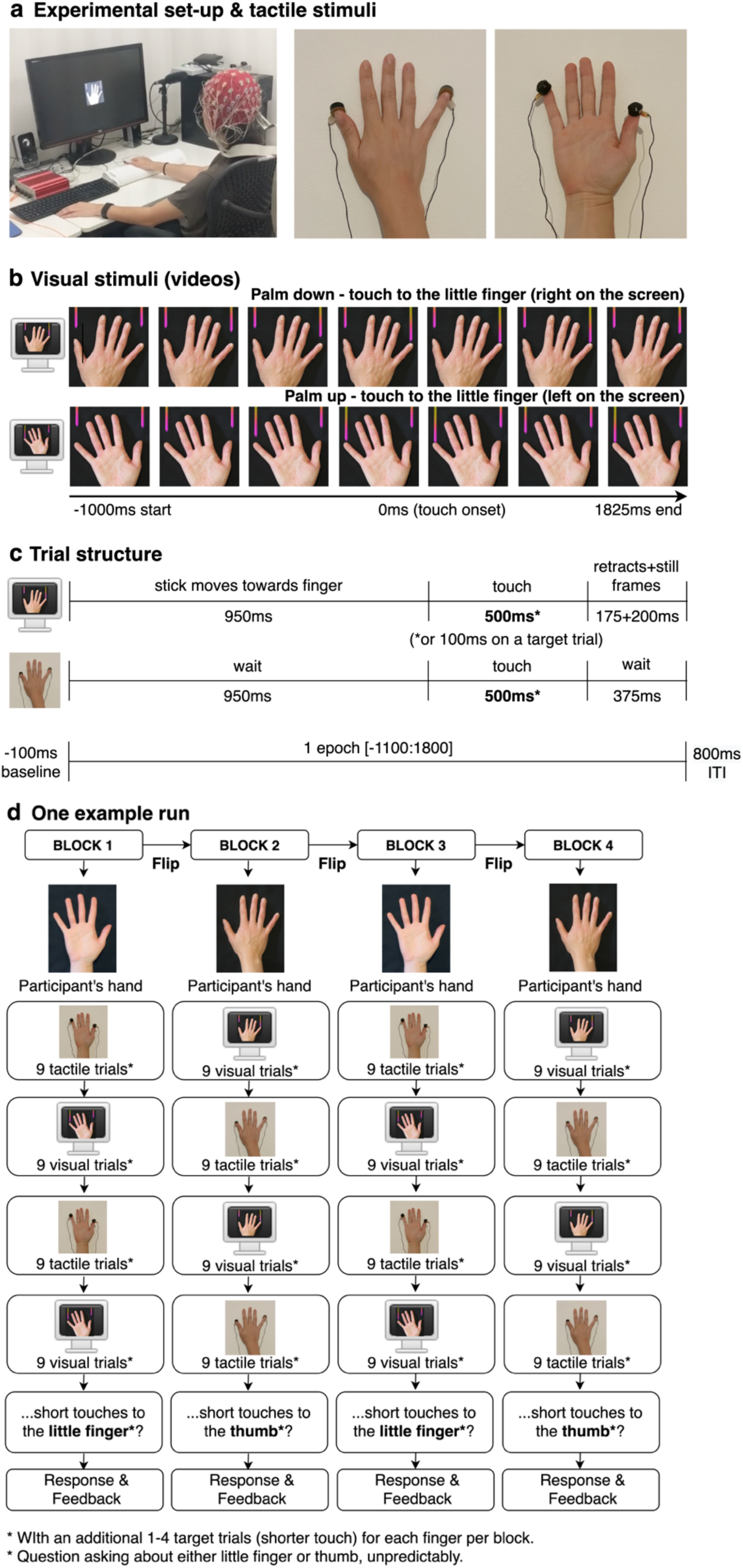
Experimental design, stimuli, and procedure. (**a, b**) Whole brain 64-channel EEG data were recorded while participants’ fingers were stimulated by two automated tactile devices (*tactile* stimuli) and while they watched videos of touch to another person’s fingers (*visual* stimuli). (**c**) Trial timing. Total trial length matched between *tactile* and *visual* trials (1825ms) with 950ms before the touch occurred and 375ms post-touch. (**d**) Each run comprised four blocks of alternating sets of nine *tactile* trials and nine *visual* trials. The orientation of the participant’s hand always aligned with the hand on the screen and both flipped between each block. Participants counted how often a target (shorter touch) occurred in each block separately for the little finger and thumb, collapsed across trial type (tactile and visual).

The visual stimuli consisted of 1825ms videos showing a single touch to either the little finger or thumb of a female right hand placed on a dark grey background. The start of each video showed an image of the same hand and two pointy metal sticks placed above the top of the little finger and thumb. As soon as the video started, one of the sticks moved towards the finger (0-950ms), touched it for 500ms (950-1450ms) before retracting (1450-1650ms) and finally remaining still (1650-1825ms; the other stick remained still throughout). We controlled the exact movement and timing of the stick by overlaying images of the stick with an image of the hand and animating it using Adobe After Effects (Christiansen, 2013). We also added shading to make the touch look realistic. We coloured the sticks with a red/pink/orange gradient to increase the saliency of the stick motion.

To ensure that the classifier could not rely on irrelevant spatial information, such as whether the stick is moving on the left or the right of the screen, the videos show the hand palm up or palm down so that half of the time a finger (little finger or thumb) was located on the left and other half on the right (Fig. 1b). We also rotated the hand slightly clockwise and anticlockwise in different videos so that the top of the little finger and thumb were either horizontally aligned or one was placed higher than the other. This resulted in 12 different videos which were shown an equal number of times throughout the experiment (videos available at: <u>doi:10.18112/openneuro.ds004563.v1.0.0</u>). On target trials we showed the same videos as described above but removed several of the frames in the middle to create a shorter touch (100ms instead of 500ms). To keep the length of the normal and target videos identical, we added frames at the end of the target videos. All videos were presented in the middle of the screen at a size of 320×320 pixels, on an AOC 27” 144Hz Monitor. Videos were presented at a sampling rate of 40 frames per second on a monitor with a refresh rate of 120Hz. Each video consisted of 73 frames which were presented in 1825ms.

The tactile stimuli consisted of a tap to the little finger or thumb of the participant’s right hand, which lasted for 500ms (normal trials) or 100ms (target trials). As with the visual trials, we added 400ms to the end of each target trial to maintain the same trial duration. The tap was produced by two electromagnetic solenoid-type vibrotactile stimulators (diameter: 18mm, height: 12mm) with a small round probe (diameter: ∼3mm, maximum extension: ∼2mm) attached with tape to the top of the fingers (Dancer Design, St. Helens, UK; dancerdesign.co.uk: Fig. 1a).

### Procedure

Participants were seated in a dimly lit room with the EEG cap fitted so that the electrode Cz was located over the vertex. The participant’s hand was placed on the table in front and aligned with the middle of the computer screen. Two small tactile devices were attached to the tip of the little finger and thumb of the right hand. To mask potential auditory noise from the tactile devices, we played white noise through speakers located on both sides of the computer screen and participants wore earplugs. After set-up was completed, participants received both verbal and written instructions and performed a test run. Once the experimenter confirmed that the task was clearly understood, the participant started the experiment by pressing the space bar.

We presented participants with alternating sets of nine consecutive touches to their own hand (*tactile* trials) and nine consecutive touch-videos on the screen (*visual* trials). Whether the run started with *tactile* or *visual* trials was counterbalanced between participants and runs. Tactile and visual stimuli were never presented at the same time (Fig. 1d). The hand orientation (palm up or palm down) in the videos changed after each block of 36 trials, and participants were instructed via a text prompt and image on the screen to flip their own hand in the same way. This ensured that the touch side for each finger always aligned between the modalities. We counterbalanced the modality, hand orientation (and thus external touch side), and touched finger within each experimental run for each participant. We also counterbalance across each run how often one trial followed on from another in order to avoid baseline issues in the decoding analysis. For example, after the little finger is touched, the next trial can be either another touch to the little finger or to the thumb (the options are little finger -> little finger, little finger -> thumb, thumb -> thumb, thumb -> little finger). We limited how often one finger could be touched in a row to a maximum of four times.

The task was for the participant to count how often a target (shorter touch) occurred in each block for the little finger and thumb separately; these trials were eliminated prior to analysis. Targets on *visual* and *tactile* trials had to be combined. The task ensured that participants were paying attention to which finger was touched but the design ensured that any task-related rehearsal or labelling strategy could not distinguish the conditions we were asking the classifier to decode. Specifically, while participants might be rehearsing (for example) “2 little finger, 1 thumb” across trials, there was no motivation to label “touch to thumb” or “touch to little finger” on each trial. There were always 1-4 targets per finger per block. After each block, a question was presented on the screen asking how many short touches the participant counted to *either* the little finger or thumb (which was probed was pseudorandom and counterbalanced; see Fig 1d). Participants answered with a number ranging from 0-9 using the numeric keypad on the keyboard and were instructed to start counting again from zero each time a new block started. Feedback about whether the response was correct or incorrect, and the right answer, was provided on the screen after each response.

There were 12 runs in total, divided into four blocks of 36 trials (with alternating sets of nine *tactile* and nine *visual* trials) resulting in a total of 1728 trials (864 tactile and 864 visual). There were an additional 240 target trials (20 per run), which were excluded from analysis. Between trials there was an inter-trial-interval (ITI) of 800ms. Each run lasted approximately 7-8 minutes with short breaks between blocks and runs. Participants controlled the length of each break and continued by pressing the space bar.

To reduce eye-movements, participants were instructed to fixate on a white fixation cross that was shown in the middle of the screen (on top of the hand during videos) throughout the entire experiment (including during *tactile* trials). However, it is important to note that eye movements are not a confounding factor interpretable by the classifier as signal, because in our experimental design the external touch side is orthogonal to the touched finger. Stimuli were presented using MATLAB (2019a, MathWorks) with Psychtoolbox (Brainard and Vision, 1997; Kleiner et al., 2007; Pelli, 1997). We used parallel port triggers to mark relevant timepoints in each trial.

After the main experiment, each participant filled out a short questionnaire consisting of three parts to assess vicarious touch experiences. First, we showed 10 videos of a ‘neutral’ (i.e., neither pleasant nor unpleasant) touch to a Caucasian female hand on a black background presented in the first-person perspective. After each video we asked participants to indicate if they felt a tactile sensation on their own body, and if so, where the touch was felt (this is a short version of our longer vicarious touch questionnaire (Smit et al., in prep). Second, we asked participants if they felt a touch on their own fingers when presented with the visual stimuli during the experiment. Third, we used a short five question Empathy Quotient scale (Muncer and Ling, 2006) to assess emotional reactivity, a dimension of empathy that relates to predicting other people’s thoughts and feelings. The entire session took approximately three hours.

### EEG data acquisition

Continuous EEG activity was recorded at a sampling rate of 2048Hz using a 64-channel Active-Two BioSemi active electrode system (BioSemi, Amsterdam, The Netherlands). This system uses a preamplifier stage on the electrode which corrects for high impedances (Metting van Rijn et al., 1990). The offset voltage was kept below 20 mV (BioSemi recommends below 50 mV). EEG electrode placement was in line with the international 10/20 standard. The EEG system was connected via a fibre optic cable to a Dell Precision T3620 computer which runs Windows 10. Data acquisition was monitored through the BioSemi acquisition software on an iMac with macOS version 10.14.1.

### Analyses

#### Vicarious touch questionnaire

Based on previous research findings (Blakemore et al., 2005; Holle et al., 2013), we anticipated that individuals who report vicarious touch sensations would have stronger neural overlap between feeling touch themselves and seeing someone else being touched, compared to those who never experience this. We therefore (as preregistered) divided our sample (n = 34) into individuals who did and did not report vicarious touch (*vicarious touch* and *no vicarious touch* groups) based on whether they reported feeling touch on their own hand (left or right) for *at least one* of the 10 videos. First, we asked: “While watching the video, did you experience a tactile sensation on YOUR body?”. If the participant answered ‘yes’ to the first question, we then asked ‘Where on YOUR body did you feel the tactile sensation?’. Only those who reported a localised touch were included in the *vicarious touch* group.

#### EEG analysis

We first downsampled the data to 256Hz using the EEGLAB toolbox (Delorme and Makeig, 2004) and re-referenced to the common average. Following a recent preprocessing method, we applied trial-masked robust detrending (van Driel et al., 2021) using the Amsterdam Decoding and Modeling toolbox (Fahrenfort et al., 2018). Like regular robust detrending (de Cheveigné and Arzounian, 2018), this method removes slow drifts in the data by first identifying glitches, which are subsequently masked out before fitting an *n^th^* order polynomial and subtracting the fit. In addition, trial-masked robust detrending also masks out cognitive events of interest, and thus the polynomial does not fit to an actual event related potential, which may otherwise result in temporal displacements or the subtraction of a real effect (for a more detailed description and the benefits over high-and low-pass filtering see (van Driel et al., 2021). We epoched and masked the data from -1100 to 1802ms (relative to the onset of touch), added a pad length of 50s, and fitted a *30^th^*order polynomial. We specified to mask current events and bad data, and to remove bad channels before fitting the polynomial. For two participants, we were unable to detrend with bad data masked. On visual inspection these datasets were normal, so we included them without the masking. We kept epochs the same length between the *tactile* and *visual* trials for easier interpretation of the data. However, whereas touch in the *visual* trials only starts at 0ms, *the onset of the visual trials* (i.e., the start of the video) is *before* 0ms (at -1000ms) and participants were able to anticipate early in the trial which finger would be touched (Fig. 1c). We applied baseline correction from -1100 to -1000 (i.e., 100ms) for both the *tactile* and *visual* trials. We did not perform any other preprocessing steps such as channel selection or artefact correction (Grootswagers et al., 2017).

#### Time-resolved multivariate pattern analysis

We applied time-resolved multivariate pattern analysis to our EEG data using MATLAB and the CoSMoMVPA toolbox (Oosterhof et al., 2016). For each participant, we extracted the *pattern of activation across all 64 EEG* channels at each timepoint and used this to train and test a linear discriminant analysis (LDA) classifier. Our classification performance metric was the average decoding accuracy across participants, for each timepoint (or timepoint combination), with 50% chance level. We performed both within-and between-modality decoding.

#### Decoding analyses of touch location (little finger versus thumb)

For the two groups separately, we tested whether neural overlap is sufficiently fine-grained such that a classifier trained on activation patterns associated with touch to the little finger versus thumb in one modality could successfully distinguish brain activation in the other modality (*between-modality* decoding). We anticipated that *between-modality* decoding might only be present for the *vicarious touch* group. However, we postulated that for both groups, a classifier should be able to extract neural signatures regarding which finger is touched *within each modality.* To verify successful within-modality decoding, we also trained and tested the classifier only on *tactile or visual* trials. We first established the decoding accuracy at each timepoint (or timepoint combination) for each participant individually and then calculated the average decoding accuracies for the two groups. Bayesian statistics were performed on the group level.

#### Between-modality decoding

It is possible that a similar pattern is elicited by seeing and feeling touch, but that this occurs at different times. To address this, we used time-generalisation methods where a classifier is trained at each timepoint on all trials from one modality (e.g., tactile) and then tested on each timepoint on all trials from the other modality (e.g., vision) (Carlson et al., 2011; King and Dehaene, 2014; Meyers et al., 2008). This method gives a time-time decoding matrix where each cell represents the average classification accuracy at that train-test timepoint combination (Fig. 2). If representations are present at the same time in both modalities, above-chance decoding will appear along the diagonal, and if they occur at different train-test timepoints, above-chance decoding will be off the diagonal.

**Figure 2.**
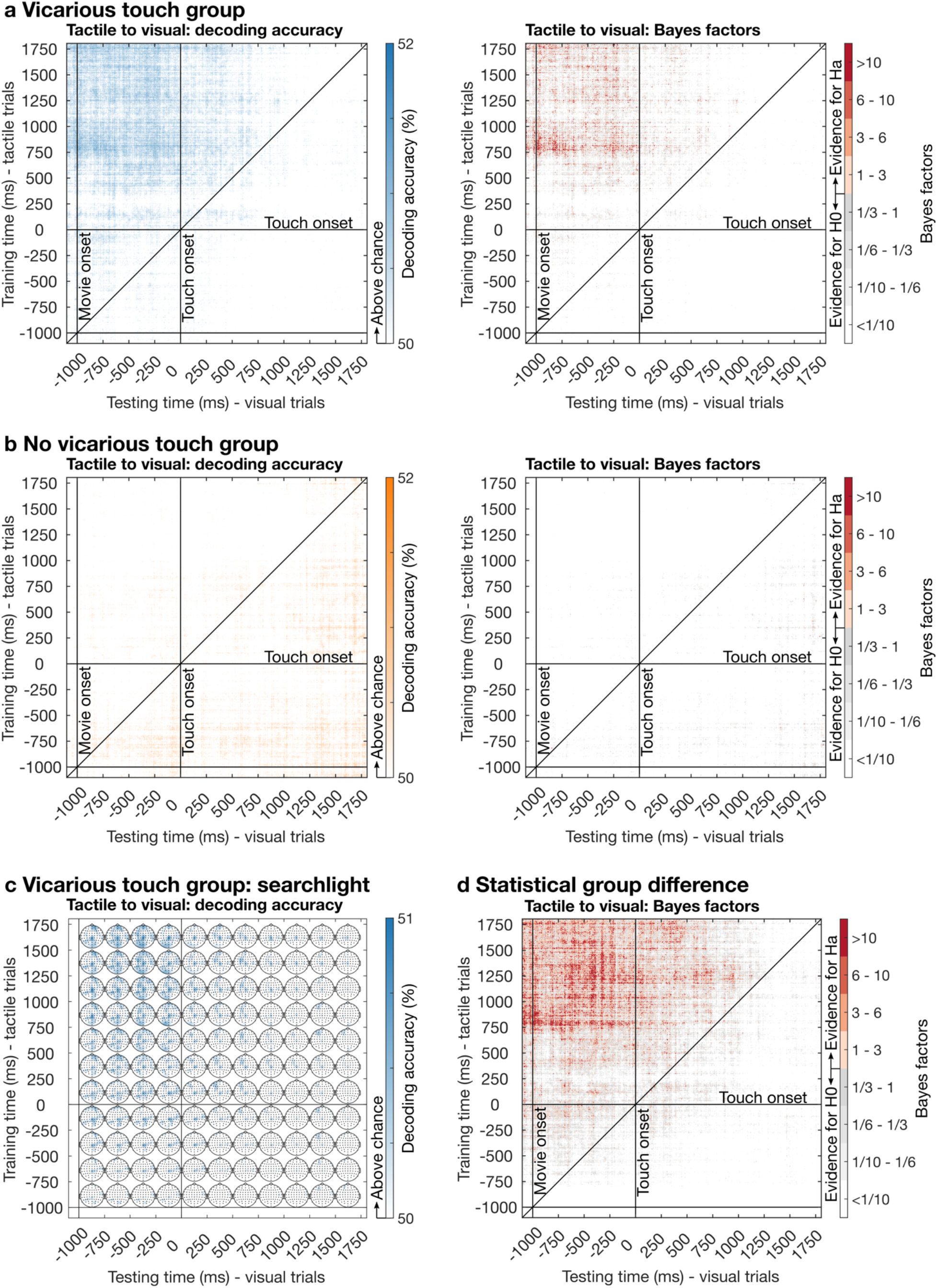
Between-modality decoding when training on *tactile* trials and testing on *visual* trials. Top and middle left plots: Mean classification accuracy for each train-test timepoint distinguishing touch to the little finger versus the thumb. Top and middle right plots: Corresponding Bayes Factors. (**a**) *Vicarious touch* group (n = 16). Above-chance decoding (>50%; blue) when training on the late *tactile* signal (∼400:1800ms) and testing on the *visual* signal (approx. -1000:750ms), with the remainder of timepoint combinations showing chance-decoding (white). BFs show the strongest evidence (red) when training on ∼750ms after touch onset in the tactile signal and between approximately -1000 and -250ms in the visual signal. (**b**) *No vicarious touch* group (n = 18). Above-chance decoding (>50%; orange) is observed sparingly, with mostly at-chance decoding (white). BFs provide very little anecdotal evidence (light orange) for between-modality decoding, with mostly strong evidence for chance-decoding (white). (**c**) Time-varying topographies showing decoding accuracies from the exploratory channel-searchlight analysis (averaged within 250ms time bins) for the *vicarious touch* group. Bayes Factors for above-chance decoding between modalities on the sensor level can be found in Supplementary Fig. 2. (**d**) Comparison at each timepoint of the *between-modality* classification accuracy between groups. BFs show the strongest evidence (red) for a difference between the groups when training on ∼750ms after touch onset in the tactile signal and between approximately -1000 and -250ms in the visual signal. There is also evidence for a difference between the groups later in the visual signal until ∼1000ms. For visualisation, all BFs >10 are plotted red, but the maximum BF is >2000.

#### Within-modality decoding

We tested whether the pattern of activation contained information regarding which finger was touched within each modality separately (combining trials from both hand orientations) at each timepoint. We used a leave-4-out cross-validation where we iteratively trained a classifier on all but four trials (one of each stimulus option, randomly selected: palm up, vs. palm down; touch to little finger vs. thumb) and tested it on the four left out trials. We only trained and tested the classifier on the same timepoints (e.g., train on 100ms and test on 100ms), as we expected that above-chance decoding would be strongest along the diagonal (where we are training and testing on the signal from the same modality).

#### Exploratory channel-searchlight (not preregistered)

To characterise information content over time and space (channels), we used an exploratory channel-searchlight analysis to identify the EEG channels that contributed to above-chance classification accuracies (see Oosterhof et al., 2016). Each EEG channel was paired with its four closest neighbouring channels to create a local cluster. We then trained and tested the classifier with these clusters (rather than whole-brain activation patterns). As with the analyses described above, for the within-modality decoding we trained and tested on the same timepoints (i.e., along the diagonal) and for the between-modality decoding we trained on *tactile* trials and tested on *visual* trials for each combination of timepoints using a time-generalisation approach. The decoding accuracy of each cluster was recorded in the central channel. We averaged decoding accuracies in the central channels over 250ms time bins (e.g., 0-250ms, 250-500ms, 500-750ms) covering almost the entire trial duration (-1000:1750ms; without 100ms baseline). We plotted the searchlight results topographically using Fieldtrip (Oostenveld et al., 2011).

#### Statistical inference

For our between-and within-modality decoding with each group, we ran Bayesian t-tests at each timepoint to establish the amount of evidence for either the alternative hypothesis (above-chance decoding; indicated by a Bayes Factor > 1) or the null hypothesis (chance decoding; indicated by a Bayes Factor < 1: Jeffreys, 1998). We used a half-Cauchy prior with medium width (r = 0.707) and adjusted the prior range of the alternative hypothesis from 0.5 to infinity to allow for small effects under the alternative hypothesis (Rouder et al., 2009). We chose this prior range because previous studies using visual decoding found effect sizes as large as 0.5 during a *baseline* period when there cannot be any meaningful information in the signal (Teichmann et al., 2021). For our *between-group comparison*, we used the same approach described above (using a half-Cauchy prior with an adjusted range) for the between-modality decoding as we expected a stronger effect for the *vicarious touch* group compared to the *no vicarious touch* group. However, for the within-modality decoding, we used a full Cauchy prior, as we did not have an a-priori hypothesis about the direction of this effect. To keep the test consistent with the directional tests, we excluded a range of d = -0.5 to d = 0.5 from the prior to exclude small effect sizes. We also performed a robustness check with a wide (r = 1) and ultrawide (*r* = 1.414) prior which did not have a pronounced effect on the results. We interpret a Bayes Factor of 6 (or 1/6 in case of evidence for the null) as substantial evidence (Jeffreys, 1998). We use a conservative criterion for accepting above-chance decoding: a single timepoint above-chance decoding may reflect noise, so we only consider evidence that classification accuracy is above chance for two consecutive timepoints based on a Bayes Factor > 6 to infer that the data at that time contain information about the location of touch. For the exploratory channel-searchlight, we calculated Bayes Factors for each channel using decoding accuracies averaged over 250ms time bins (Supplementary Fig. 2 and 3). For these Bayesian analyses we used the Bayes Factor R package (Morey et al., 2018).

**Figure 3.**
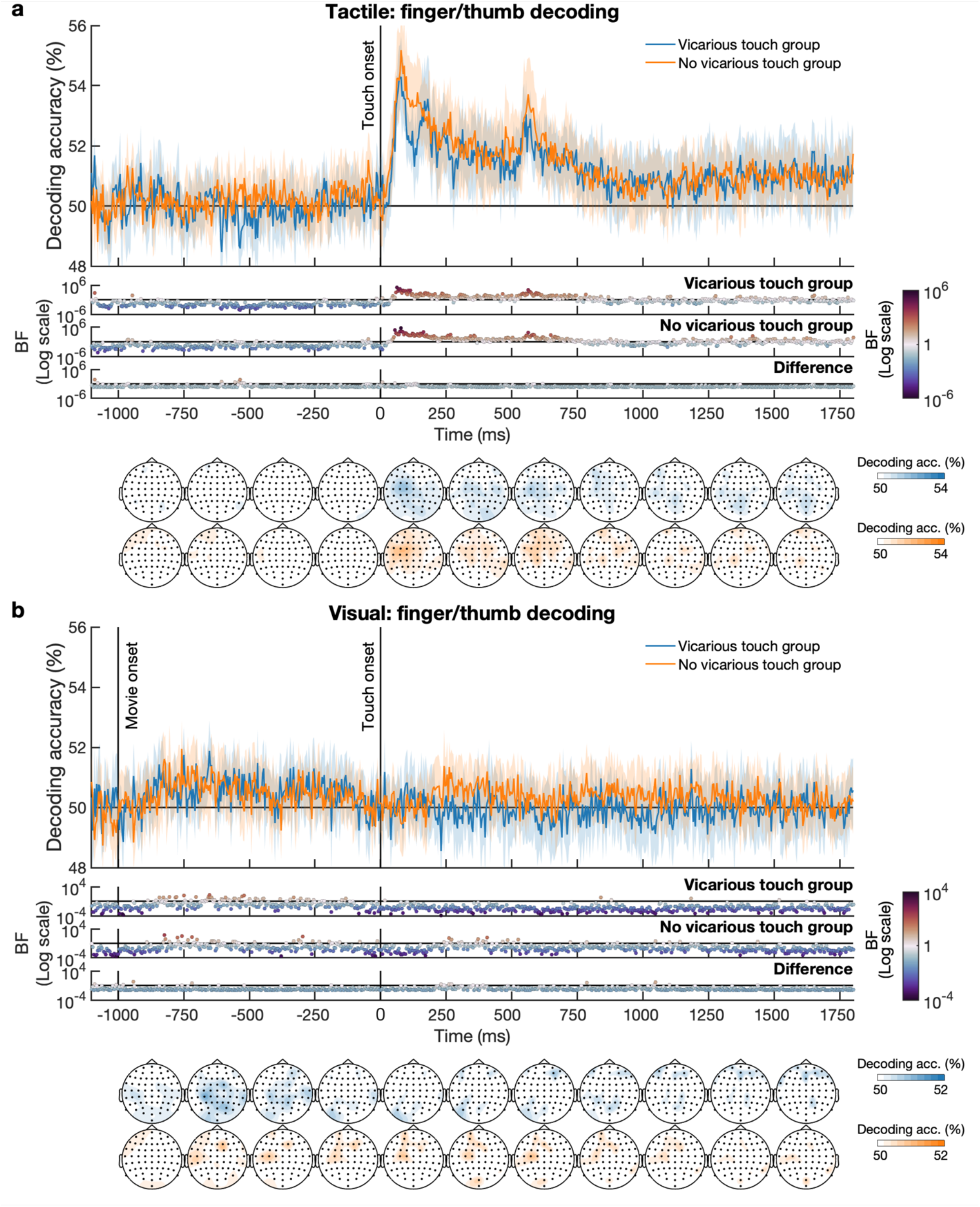
Within-modality decoding for the tactile (top) and visual (bottom) modality. Mean classification accuracy over time for the *vicarious touch* group (n = 16; blue) and the *no vicarious touch* group (n = 18; orange). In all plots, shading indicates 95% confidence intervals. Dots below plots show Bayes Factors for each time point, indicating whether there is evidence for the alternative hypothesis (red: above-chance decoding) or the null (blue: chance decoding); note the different Bayes Factor scales for top and bottom panels. Time-varying topographies showing decoding accuracies from the exploratory channel-searchlight analysis (averaged within 250ms time bins) are displayed below for each modality and group. Bayes Factors for above-chance decoding within each modality on the sensor level can be found in Supplementary Fig. 3. (**a**) Tactile decoding plots (top) show an initial baseline period of 1100ms (100ms baseline + 1000ms wait; see Fig. 1) and touch occurs from 0-500ms. The high decoding peaks occur at the onset and offset of the touch. For both groups, the location of touch can be decoded for *tactile* trials from ∼45ms after touch onset. (**b**) Visual decoding plots (bottom) show an initial baseline period of 100ms followed by video onset at -1000ms and touch occurs between 0-500ms. For the *vicarious touch* group, the location of touch can be decoded for the *visual* trials during the anticipatory period between ∼170ms after the start of the video until ∼130ms before the touch happened. For the *no vicarious touch* group, above-chance decoding was present in the *visual* trials from ∼160ms after the start of the video and again later in the trial between ∼260-400ms during the touch (touch is 0-500ms). Bayes Factors provide evidence for no difference between the groups (bottom).

For the emotional reactivity empathy scale, we use a Bayesian Mann-Whitney Independent Samples t-test to assess whether the *vicarious touch* group scored higher than those in the *no vicarious touch* group (using the statistical software JASP, Version 0.17.1, 2023).

#### Preregistration

We preregistered this study, including a detailed analysis plan, on the Open Science Framework (OSF) before data collection. Prior to running analyses, we decided to make the following changes to the preregistered analysis methods. We used trial masked robust detrending instead of applying a high-and low-pass filter as this has recently (after preregistration) been shown to introduce fewer artifacts, especially for time generalisation (van Driel et al., 2021). For the within-modality decoding, we modified the folds as the EEG task lasted for approximately two hours and participants reported that it was demanding, making it likely that physiological artifacts (e.g., blinking, moving, sweating) and cognitive artifacts (e.g., tiredness, lack of concentration) would increase throughout the task. Initially we planned to use a twelvefold cross-validation where we train the classifier iteratively on 11 runs and test it on the left out run, but this could be influenced by any drift or change across the experiment. Instead, we used a leave-4-out approach, where we iteratively trained on all but four trials (one of each stimulus option: palm up, vs. palm down; touch to little finger vs. thumb) and tested it on the four left out trials. This way, we train the classifier on trials from the beginning, middle and end of the experimental task. We also applied a baseline correction for the within-modality analysis^2^. Using baseline correction before decoding has been shown to increase the likelihood of picking up on an effect without introducing artefacts (at least when trial-masked robust detrending is used rather than high-pass filtering: van Driel et al., 2021), potentially by reducing temporal drifts due to internal and external sources (Tanner et al., 2016). We had preregistered to analyse the groups separately but not specifically the between-group statistical comparison, which we conducted to assess whether the *vicarious touch* group was statistically different from the *no vicarious touch* group (as opposed to just being able to state that they show an effect, whereas the *no vicarious touch* group showed evidence for no effect). Finally, in response to reviewer comments, we included an exploratory channel-searchlight analysis.

### Data availability

The raw and preprocessed data, analysis and experiment presentation scripts, and stimuli are available in BIDS format via OpenNeuro: <u>doi:10.18112/openneuro.ds004563.v1.0.0</u>.

## Results

### Vicarious touch questionnaire

Our final sample after exclusions consisted of 16 participants classified as having vicarious touch and 18 participants as having no vicarious touch based on their responses to the vicarious touch video questionnaire (Smit et al., in prep). Of the 17 participants who answered ‘yes’ to “While watching the video, did you experience a tactile sensation on YOUR body?”, 16 indicated the sensation was localised only to the left hand (n = 8), only to the right hand (n = 4) or to alternating/both hands (n = 4). One participant indicated that the mirrored sensation was ‘a general feeling’ and we therefore placed this participant in the *no vicarious touch* group (n = 18). For the *vicarious touch* group, the average number of times a participant reported a tactile sensation when watching the videos depicting touch was five (see Table 1). Six out of 16 participants had scores >5 which could indicate mirror-touch synaesthesia based on previous definitions (Ward et al., 2018). However, we followed up with these six participants and none reported vicarious tactile sensations outside the experimental setting, inconsistent with mirror-touch synaesthesia (which has an estimated prevalence of only 1.6%; Banissy et al., 2009). Three participants belonging to the *vicarious touch* group answered ‘yes’ to the question: “During the experiment you just participated in, did you feel a touch on your own body when you saw the metal stick in the video touch the hand on the screen?” One participant wrote the following description: “I felt the touch on my right hand, sometimes on my thumb and sometimes on my pinky depending on where the metal stick touched the hand on the screen.” None of the participants in the *no vicarious touch* group answered ‘yes’ to this question.

For the emotional reactivity empathy scale, we used a Bayesian Mann-Whitney Independent Samples t-test to assess whether the *vicarious touch* group scored higher than those in the *no vicarious touch* group. This provided evidence for the null hypothesis (BF = 0.126; no greater emotional reactivity).

### Between-modality decoding of touch location (little finger versus thumb)

In the *vicarious touch* group, there was strong evidence that information about the location of touch (little finger or thumb) was present in the neural activation patterns evoked when seeing or feeling touch, and that these patterns overlapped between modalities (Fig 2a). As a group, the average between-modality decoding accuracy was 52%, with the maximum individual decoding accuracy being 56%. The statistics show that these numerically small classification accuracies are robust and demonstrate that the classifier can reliably categorise touch location above-chance when trained on *tactile* and tested on *visual* trials. The timecourse of this overlap is key: the neural pattern 400ms onwards after directly feeling touch overlaps with a much earlier time when seeing touch - consistent with a prediction of upcoming touch location. The strongest evidence is present between the late tactile signal (∼750ms after touch onset) and the anticipatory part of the visual signal (video start until ∼250ms before touch onset). This suggests that we are not seeing initial early somatosensory processing but rather a more abstracted touch representation activated in these individuals when predicting upcoming touch locations in the videos. The *no vicarious touch* group showed almost exclusively evidence for chance-decoding (Fig. 2b). We also trained the classifier on *visual* trials and tested on *tactile* trials, which produced similar but weaker results (see Supplementary Fig. 1). This asymmetry in the direction of generalisation is likely due to variation in signal-to-noise between datasets (Hurk and Beeck, 2019), and may reflect the greater temporal uncertainty in the videos which increases the trial-by-trial and between-participant variability in the visual domain. The exploratory channel-searchlight analysis for the *vicarious touch* group suggests that between-modality decoding of touch location is driven by a distribution of channels across the head, including what appears to be the somatosensory cortex (Fig. 2c; for corresponding channel-searchlight statistics see Supplementary Fig. 2). The limitations for interpreting spatial effects are addressed in the discussion.

We also tested directly if the *vicarious touch* group had higher decoding accuracies than the *no vicarious touch* group at each timepoint (post hoc analysis; one-tailed Bayesian t-tests). This demonstrated strong evidence in terms of the degree to which between-modality decoding was present (Fig. 2d). This is an additional statistical verification of the presence of *between-modality* decoding in the *vicarious touch* group and the lack of such decoding in the *no vicarious touch* group.

### Within-modality decoding of touch location (little finger versus thumb)

To assess whether a classifier can successfully extract neural signatures distinguishing touch to the little finger versus thumb *within each modality*, and at what time, we trained and tested a classifier using only the tactile *or* visual data. For *tactile* stimuli, the classifier was able to predict above chance (BF >10) which finger was touched from ∼45ms after touch onset for both groups, and this was maintained throughout the trial (Fig. 3a). This shows the remarkable sensitivity of these methods, considering the close proximity of finger representations in somatosensory cortex (Kuehn et al., 2014). For *visual* stimuli, decoding of touch location was also present but much more subtle, reflecting the indirect information about touch (watching a video) and temporal variability when making inferences about an unfolding touch event. For visual stimuli, the *vicarious touch* group showed some above-chance decoding during the anticipatory period in the trial (∼170ms after video start until ∼130ms before the touch onset; Fig. 3b) suggesting information about the perceived location of touch on another person’s body is maximally present when anticipating where the touch will occur, rather than during the touch. The *no vicarious touch* group showed some above-chance decoding for a short period ∼160ms after video onset, and again later in the trial between ∼260-400ms during the touch (Fig. 3b). However, when we directly tested for a difference between the groups (post hoc analysis; two-tailed Bayesian t-tests), there was evidence for no difference in decoding accuracy between the groups in either the visual or tactile domains (Fig. 2a, b). The exploratory channel-searchlight analysis for both groups suggests that within-modality decoding of touch location is driven by a distribution of channels across the head (Fig. 3a,b; for corresponding channel-searchlight statistics see Supplementary Fig. 3).

## Discussion

The aim of this preregistered study was to enhance our understanding of the neural mechanisms underlying vicarious touch. We used time-resolved MVPA of whole-brain EEG data recorded while participants directly felt touch to different fingers or observed similar touch to another person. We examined *if* there is neural overlap between seeing and feeling touch, and if so, *when* this is present. This provides insight into shared touch representations and if these are of an early (sensory) or later, more abstract, processing stage. Our main finding was that only in people with vicarious touch experiences (who reported feeling touch when they watched videos of touch) did neural activation patterns overlap *between* modalities, and this contained information about the location of touch. The chance-level decoding between modalities for the group of people who did not report such vicarious touch experiences, and the strong evidence for a difference between the groups, provide a link between overlapping neural representations and subjective experiences. In the *vicarious touch* group, late tactile patterns overlapped with anticipatory visual patterns: seeing touch activated information that occurred 400ms after being touched directly – therefore presumably reflecting more abstracted information. The *within-modality* tactile decoding results demonstrate that information about which finger was touched is already present very early on (∼45ms onwards) for both groups, but this does *not* seem to be what is instantiated when we see touch.

Vicarious tactile experiences could rely on a simulation process whereby seen touch is mapped onto a similar representation within the observer’s own system. Some simulation theories propose that we empathise with other people’s sensations by first representing the observed event in our own relevant perceptual areas. This representation is then ‘directly matched’ against sensations we could experience first-hand. Such an automatic process allows for passive ‘resonating’ with another person (e.g., Iacoboni et al., 1999; Rizzolatti et al., 2001; Rizzolatti and Sinigaglia, 2016). Other simulation theories argue that tactile empathy could be better explained as a prediction process (for a review see Bach and Schenke, 2017). Such ‘predictive coding’ accounts propose that social interactions, including vicarious touch, are hypothesis driven, such that an assumption about the external world is transformed into predictions about the expected perceptual consequences of the observed event. Prediction errors about an observed touch event without direct tactile input can be ‘explained away’ by activity in the somatosensory cortex (Bach and Schenke, 2017; Ishida et al., 2015). Both these accounts propose that seeing touch should evoke corresponding *sensory* representations (see also Banissy and Ward, 2007; Bufalari and Ionta, 2013; Keysers and Gazzola, 2009; Schaefer et al., 2009). This predicts that the information that is present early on when we are touched should also be instantiated when we see someone else being touched. In our experiment, this would mean that *between-modalit*y decoding should involve early tactile information appearing within the visual patterns.

Certain aspects of our *between-modality* decoding results are consistent with simulation theories. Tactile information did appear in *visual* trials before the stick touched the hand in our *vicarious touch* group. This suggests that this information is involved in predicting the consequences of an upcoming touch. When the touch actually occurred in the video, the *between-modality* decoding dropped to chance. This is consistent with a predictive coding account whereby the lack of predicted tactile stimulation then modulates the neural response. However, our Bayesian analysis provides strong evidence that the early tactile representations (<400ms after tactile stimulation) *do not* generalise to the visual representation. This indicates that any simulation or prediction of seen touch operates at a higher or more abstract level than typically proposed (see also Kilteni et al., 2021). Previous univariate EEG studies found both early (45ms) and later (up to 700ms) somatosensory modulations when participants saw touch, which has been taken as evidence that shared representations are already present at very early processing stages (Adler et al., 2016; Adler and Gillmeister, 2019; Bufalari et al., 2007; Martínez-Jauand et al., 2012; Rigato et al., 2019b, 2019a; Streltsova and McCleery, 2014). However, most of these EEG studies test how seeing touch modulates somatosensory evoked potentials – and having the seen and felt touch simultaneously presented means it is difficult to disentangle multisensory integration modulations from shared neural representations. Our results suggest that only the later effects are likely to reflect shared touch representations, and that this would depend on whether the participant tends to generally experience vicarious touch or not. Our findings have important implications for simulation theories as the data suggest that vicarious touch does not rely on the activation of a low-level sensory representation of the touch.

Within the framework of simulation theories and the idea that overlapping neural representations might be a mechanism for tactile empathy (Bufalari and Ionta, 2013; Keysers and Gazzola, 2009), our *vicarious touch* group could be classified as having ‘high tactile empathy’ and our *no vicarious touch* group as ‘low tactile empathy’. We did not find higher scores for emotional reactivity (the tendency to react emotionally to others’ feelings) in the *vicarious touch* group, which suggests that any increased empathising with others might be specific to sharing bodily sensations, rather than a general higher emotionality. This differs from other studies that found higher emotional reactivity scores in mirror-touch synaesthetes compared to controls (Banissy and Ward, 2007; Ward et al., 2018).

Although tactile empathy is one possible explanation for our group differences, there are also other potential contributors. For example, it has been proposed that the phenomenon of somatosensory activity evoked by the mere sight of touch is rooted in the inherently multi-modal nature of our perception of the body (de Vignemont, 2017). This perspective posits that visual information about body parts, regardless of whether they belong to the self or others, is automatically integrated with tactile information without vicarious touch providing any additional information about the other person. Further, as touch was presented to a hand in a first-person perspective, it is possible that the observed effects stem from own-body perception rather than (or in addition to) empathetic responses to another person’s sensations (see also Adler and Gillmeister, 2019). Viewing an object near the hand could also involve peripersonal space mechanisms. This involves visual-tactile (bi-modal) neurons with tactile receptive fields anchored to a specific body part that can also respond to visual stimuli near the observer’s own body part and sometimes to stimuli close to the corresponding body part of another person (Brozzoli et al., 2013; Graziano and Cooke, 2006; Ishida et al., 2010). Such peripersonal space mechanisms located in parietal and frontal brain areas could be involved in the shared encoding of the visual stimulus *approaching the finger* and the *touch on the finger*, which is consistent with the timecourse of overlap in our study. Finally, individuals with vicarious touch tendencies could generally have a more organised internal representation of directly experienced touch. This would provide the classifier with a less noisy dataset for training on *tactile* trials, potentially leading to improved generalisation on *visual* trials. In our data, however, both groups show high decoding accuracy for the tactile stimuli with evidence for no difference between the groups (Fig. 3a), making this seem less likely. It is always possible though that with more fine-grained tactile stimulation, where the touches are presented closer to each other on the body, we might see evidence of a noisier representation in those who do not experience vicarious touch. Future studies could test these alternative/additional mechanisms by presenting touch closer on the body (e.g., the middle and ring finger) or by showing touch to a hand in an other-specifying (allocentric) orientation and/or outside the participant’s peripersonal space.

The focus of our study was on testing the timecourse of any shared neural representations using time-resolved multivariate decoding methods which rely on the *pattern of activation across many sensors* (in our case 64 channels across the whole brain). As such, inferences about which individual areas of the brain contribute to the above-chance classification can only be speculative. An exploratory channel-searchlight analysis on the within-modality decoding suggests that channels located over the primary and/or secondary somatosensory cortex contribute to above-chance decoding in the tactile domain (Fig. 3), which would be expected. For the between-modality decoding, a distribution of channels located across the brain contributed to above-chance decoding (Fig. 2). However, as EEG has high temporal but low spatial resolution, it is better suited for answering questions about *when* rather than *where*. As such, we do not know if the somatosensory cortex is predominantly driving the observed neural overlap between seeing and feeling touch. Indeed, it may well be an amodal representation of touch that is shared between the modalities. For instance, regarding vicarious pain perception, one study using voxel-based morphometry found that participants who reported vicarious pain showed increased grey matter volume in the primary somatosensory cortex, compared to those who did not report such experiences (Grice-Jackson et al., 2017). However, another study using MPVA/fMRI found that different areas computed somatotopy (pain to upper versus lower limbs) for somatic pain (somatosensory cortex) and vicarious pain (mentalising-related circuits) (Krishnan et al., 2016). As such, the role of the somatosensory cortex in vicarious sensory processing remains unclear. Our study was focused on the timecourse of any overlap, which clearly suggests a distinction between the rapid somatosensory processing involved when directly feeling a tactile stimulus (∼45ms) (Allison et al., 1992) and more extensive processing involved when seeing touch (>400ms). This tells us that visually perceiving touch evokes a more abstracted representation of directly felt touch, rather than an early sensory representation.

Our findings provide a link between the overlap in neural activation patterns evoked by seeing and feeling touch and the subjective experience of vicarious touch. EEG is correlational by nature, and therefore, while the results provide insight into neural overlap between seen and felt touch co-occurring with vicarious touch experiences, we cannot say whether the shared representations are *necessary* for differences in phenomenology – they could be epiphenomenal (e.g., de-Wit et al., 2016; Lamm and Majdandžić, 2015). Further, chance-decoding does not guarantee that the information is not present in the brain, it may simply be that MPVA and EEG are unable to detect it (de-Wit et al., 2016). Another issue that often arises in the context of multivariate decoding is whether one is using it for *interpretation* or trying to *maximise prediction*. Here, we are testing the evidence that a neural pattern instantiated when feeling touch is also present when viewing touch: using decoding for interpretation. Thus, *any* reliable decoding above chance shows there is structure in the data that allows for categorical distinction between conditions (Hebart and Baker, 2018). Our *between-modality* decoding accuracies are low (∼52%) but this is not indicative of effect size (Combrisson and Jerbi, 2015; Hebart and Baker, 2018). It is important to note that we use minimal preprocessing (e.g., no filtering, trial rejection, trial averaging, channel-selection) to ensure that our results do not rely on specific preprocessing choices (Grootswagers et al., 2017). However, this also means that the data are noisier and can result in lower decoding accuracies. Additionally, the careful control of the video stimuli to eliminate external touch signals from driving the classifier, coupled with potential temporal variability between trials and participants in observing the visually unfolding touch event, make it unlikely for high decoding accuracies to be achieved in the *visual* trials. Finally, the brain’s response to feeling touch to the fingers and seeing videos of touch presumably should be quite distinct – we are presenting completely different stimulation of different modalities. Despite these extensive sensory differences, the cross-generalisation analysis can still predict touch location across modalities, showing there is (at some level) information about touch location that is activated when some people see touch. The Bayesian evidence demonstrates the reliability of this overlap. With the above caveats in mind, our findings provide new evidence suggesting that shared touch representation could be a potential mechanism underlying vicarious touch.

Another aspect of our methodology that is crucial to note is that a classifier can pick up not only the effect of interest (here, which finger was touched), but also any other feature that differs between the conditions of interest (e.g., mental labelling, low-level stimulus features, systematic eye movements: Alizadeh et al., 2017; Woolgar et al., 2014). If participants were mentally labelling trials as “little finger” or “thumb”, for example, this could be an alternative explanation for our *between-modality* decoding. In our task, however, labelling each trial as such would interfere with completing the task (cumulative counting of targets to each finger across trials), and the target trials were excluded from analyses. Further, it is unclear why any such labelling strategy would differ between people with and without self-reported vicarious touch sensations. Our careful experimental design also controls for external touch side, low-level visual features and systematic eye-movements (by having touch side orthogonal to the decoded feature of which finger was touched). This, in combination with strong evidence for shared representations only in the *vicarious touch* group, supports the interpretation that our *between-modality* decoding reflects overlap in neural patterns that are *specific to the manipulated variable of interest (i.e., touch to the little finger versus thumb).* In this study, we did not measure how *vicarious touch* participants felt the touch in the videos relative to the touch they actually received; future work could look at whether a closer match between the seen and felt touch increases between-modality decoding.

Individuals who reported vicarious touch could also be involuntarily altering their phenomenology to meet implicit task requirements, potentially indicating a trait difference in their ability to control their perceptual experiences (Lush et al., 2020). This idea is closely linked to the possibility that imagery may serve as the mechanism for vicarious experiences (e.g., Jacob and de Vignemont, 2016). To further explore this idea, future studies could instruct participants to actively imagine the touch being applied to their own bodies when they see it, and then evaluate the impact on the presence and timecourse of neural overlap (for a similar approach with vicarious pain see Krishnan et al., 2016). In addition, previous vicarious pain studies report sex differences (Singer et al., 2006; Yang et al., 2009), and this may also have had an influence in the current study as our sample consisted of more females (68%; however, the men in the study did fall into both groups; four in the *vicarious touch* group and seven in the *no vicarious touch* group).

The presence of overlapping neural representations at an anticipatory visual and late tactile processing stage might explain why most individuals report vicarious touch only in certain contexts, such as when specifically attending to experimental videos of touch. In contrast, mirror-touch synaesthetes feel other people’s sensations in non-experimental contexts, which might indicate more pronounced ‘tactile empathy’ (e.g., Blakemore et al., 2005; Keysers et al., 2010; Ward et al., 2018; but see de Vignemont, 2017 challenging this). It is therefore possible that for mirror-touch synaesthetes, seeing someone else being touched evokes comparatively earlier (more low-level or sensory) tactile representations. Our experimental paradigm allows future investigation into the temporal characteristics of neural representations in mirror-touch synaesthesia and neurodiverse populations in which self-other processing is altered (e.g., psychopathy, autism, alexithymia or depersonalisation disorder: Bird and Viding, 2014; Gillmeister et al., 2017).

In conclusion, our results increase the understanding of how vicarious touch is processed in the brain by showing that seeing another person being touched can activate representations overlapping with those evoked by feeling touch directly. Such shared representations were only observed in people who reported feeling touch when seeing videos of touch, suggesting a link between shared representations and the subjective experience of vicarious touch. The timecourse of our findings provide insight into the underlying mechanism for vicarious touch, as it implies that observing an approaching touch to another person evokes an abstracted representation of directly felt touch, rather than the early somatosensory representation typically proposed by simulation theories.

## Author Contributions (CRediT)

**SS**: Conceptualization, Methodology, Software, Investigation, Data Curation, Formal analysis, Visualization, Writing – Original Draft, Writing – Review & Editing, Project administration. **DM**: Conceptualization, Methodology, Software, Formal analysis, Visualization, Writing – Review & Editing. **RZ**: Conceptualization, Writing – Review & Editing, Supervision, **ANR**: Conceptualisation, Methodology, Writing – Review & Editing, Supervision.

## Acknowledgments

We thank Alexandra Woolgar, Chris Baker and Simmy Poonian for valuable feedback on the manuscript.

## Footnotes

1 We use the term “information” to refer to a statistical dependence in the data that can be “read out” using machine learning methods (Hebart and Baker, 2018), but see de-Wit et al., (2016) for important limitations on whether this this information is used by the brain.

2 We established within-modality decoding accuracies for the first four participants (without running any group statistics) before realising that baseline correction was necessary to reduce temporal drifts due to the length of the experiment, so we applied this to the entire sample prior to statistical analysis.

