## Supplementary Materials for "Vicarious touch: overlapping neural patterns between seeing and feeling touch"

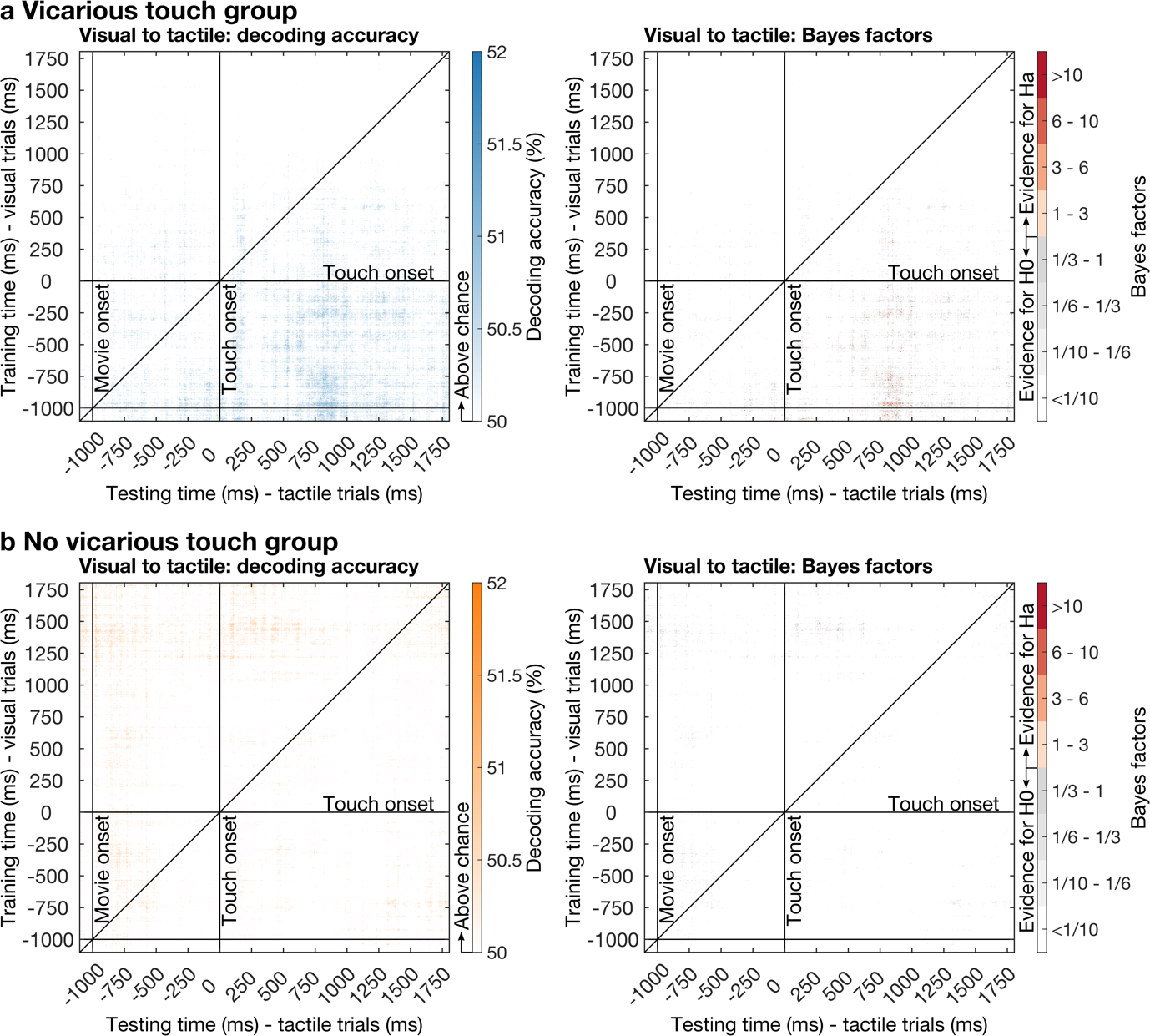

**Supplementary Figure 1:** **Between-modality decoding when training on *visual* trials and testing on *tactile* trials.** Left plots: Mean classification accuracy for each train-test timepoint distinguishing touch to the little finger versus the thumb for the (**a**) *Vicarious touch* group (>50%; blue, n = 16) and (**b**) *No vicarious touch* group (>50%; orange, n = 18). Right plots: Corresponding Bayes Factors showing predominantly strong evidence for the null (chance-decoding; white) with some moderate-strong evidence for the alternative (above-chance decoding; orange/red) in the *vicarious touch* group.

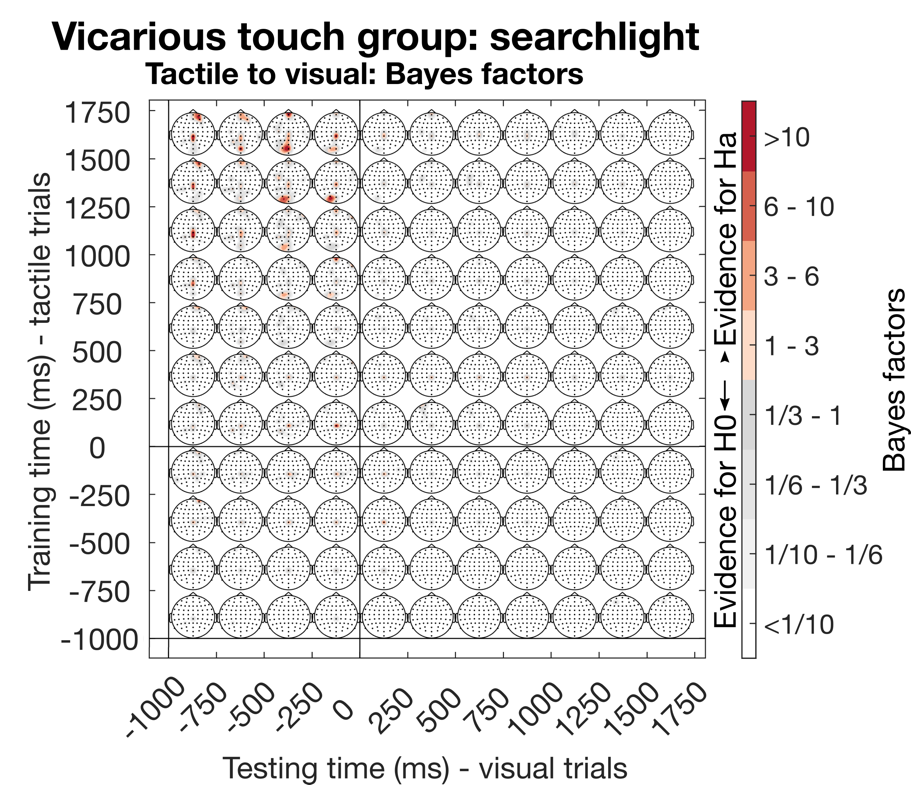

**Supplementary Figure 2: Between-modality exploratory channel-searchlight statistics.** Time-varying topographies showing Bayes Factors from the exploratory channel-searchlight analysis for the *vicarious touch* group when training on *tactile* trials and testing on *visual* trials (decoding accuracies are presented in the main paper). Statistics are calculated using averaged decoding accuracies over 250ms time bins. Bayes Factors indicate evidence for above-chance decoding on the sensor level (orange/red) with white indicting evidence for chance-decoding.

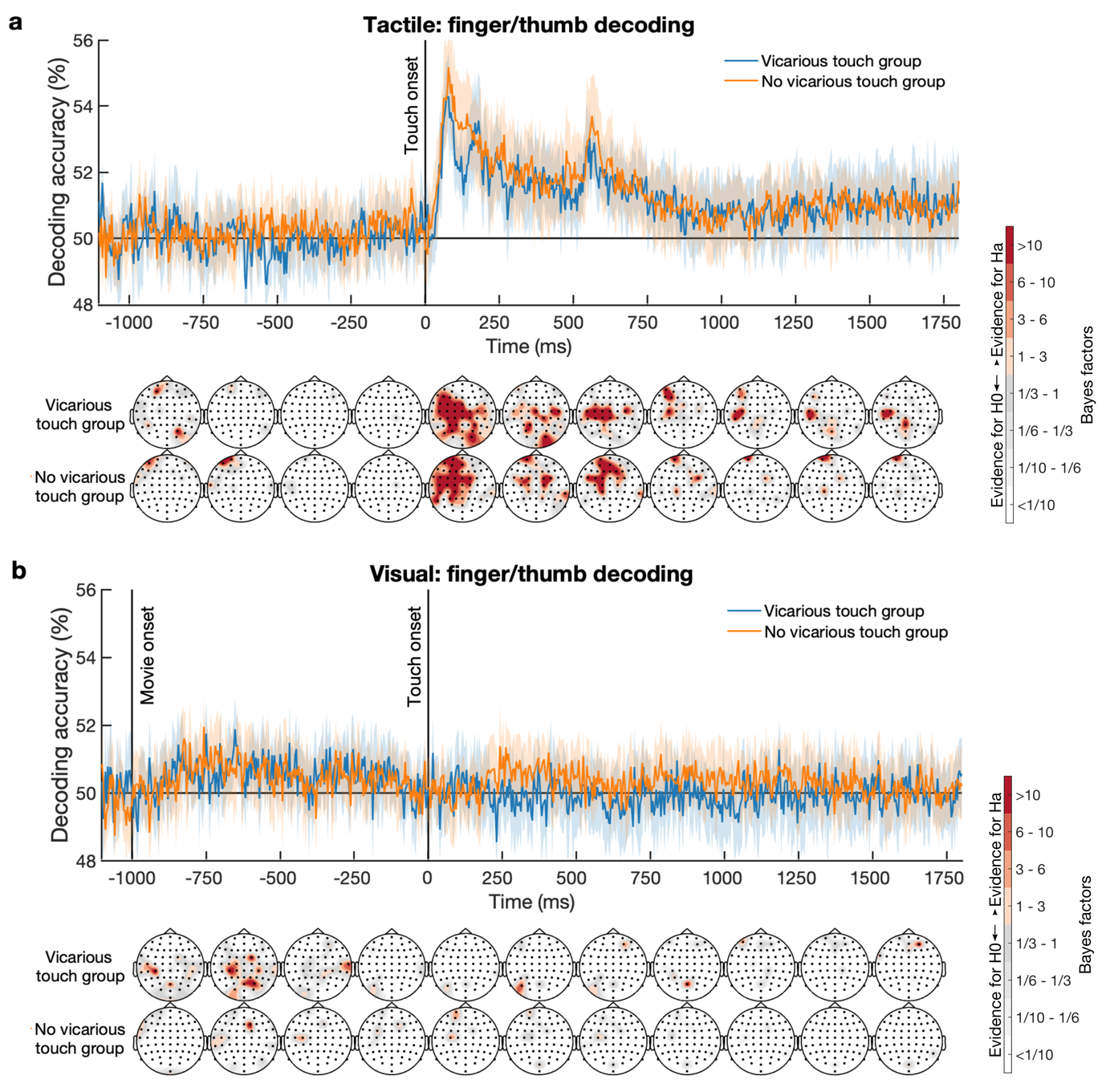

**Supplementary Figure 3:** **Within-modality exploratory channel-searchlight statistics.** Mean classification accuracy over time for the *vicarious touch* group (n = 16; blue) and the *no vicarious touch* group (n = 18; orange). In all plots, shading indicates 95% confidence intervals. Time-varying topographies showing Bayes Factors from the exploratory channel-searchlight analysis are displayed below for each modality and group (decoding accuracies are presented in the main paper). Statistics are calculated using averaged decoding accuracies over 250ms time bins. Bayes Factors indicate evidence for above-chance decoding on the sensor level (orange/red) with white indicting evidence for chance-decoding.
